# Structural O-Glycoform Heterogeneity of the SARS-CoV-2 Spike Protein Receptor-Binding Domain Revealed by Native Top-Down Mass Spectrometry

**DOI:** 10.1101/2021.02.28.433291

**Authors:** David S. Roberts, Morgan W. Mann, Jake A. Melby, Eli J. Larson, Yanlong Zhu, Allan R. Brasier, Song Jin, Ying Ge

## Abstract

Severe acute respiratory syndrome coronavirus 2 (SARS-CoV-2) utilizes an extensively glycosylated surface spike (S) protein to mediate host cell entry and the S protein glycosylation is strongly implicated in altering viral binding/function and infectivity. However, the structures and relative abundance of the new O-glycans found on the S protein regional-binding domain (S-RBD) remain cryptic because of the challenges in intact glycoform analysis. Here, we report the complete structural characterization of intact O-glycan proteoforms using native top-down mass spectrometry (MS). By combining trapped ion mobility spectrometry (TIMS), which can separate the protein conformers of S-RBD and analyze their gas phase structural variants, with ultrahigh-resolution Fourier transform ion cyclotron resonance (FTICR) MS analysis, the O-glycoforms of the S-RBD are comprehensively characterized, so that seven O-glycoforms and their relative molecular abundance are structurally elucidated for the first time. These findings demonstrate that native top-down MS can provide a high-resolution proteoform-resolved mapping of diverse O-glycoforms of the S glycoprotein, which lays a strong molecular foundation to uncover the functional roles of their O-glycans. This proteoform-resolved approach can be applied to reveal the structural O-glycoform heterogeneity of emergent SARS-CoV-2 S-RBD variants, as well as other O-glycoproteins in general.

## Introduction

The novel zoonotic 2019 coronavirus disease (COVID-19) global pandemic^1^ has led to more than 102 million reported cases and over 2 million deaths as of February 2021.^2^ The causative pathogen of COVID-19, severe acute respiratory syndrome coronavirus-2 (SARS-CoV-2), utilizes an extensively glycosylated spike (S) protein that protrudes from the viral surface to bind receptor angiotensin-converting enzyme 2 (ACE2) for cell entry.^3,4^ The evolution of the surface S protein is particularly important as it initiates pathogenesis,^5^ is the main target of vaccination^6,7^ and antibody therapeutic design,^8,9^ and a substantial number of predicted mutations in the S protein regional-binding domain (RBD) can potentially enhance ACE2 binding.^10^ Moreover, the more than 636,000 viral genomic sequences generated from global molecular epidemiology studies (as of February 28, 2021, www.gisaid.org)^11^ reveal the emergence of a number of variants in the S protein, indicating that SARS-CoV-2 may be evolving to acquire selective replication advantages, drug insensitivity and immunological resistance, thereby underscoring the need for new technologies capable of rapid and deep structural profiling of the virus.^12-14^

Importantly, glycans flank the polybasic furin cleavage site of S protein necessary for S stability during biogenesis and influence conformational dynamics by masking protein regions from cleavage during ACE2 binding.^3,15,16^ How these different sites are glycosylated and how they influence ACE2 binding are likely to affect cell infectivity and could shield certain epitopes from antibody neutralization.^17-20^ The SARS-CoV-2 S protein carries 22 N-glycosylation sequons per protomer which have been characterized in detail.^21,22^ However, characterization of O-glycosylation has so far been limited, with only putative O-glycosites reported,^16,22,23^ and the glycan microhetereogeneity, including molecular compositions, structures, and relative abundance, remains cryptic due to the challenges in O-glycan analysis.^5,17,24-29^ Importantly, individual glycosites can give rise to many glycan structural variants, *i*.*e*. glycoforms, and these differences in glycan structures, even at a single glycosite, can have critical implications on biological functions.^20,30-32^ It is crucial to decipher the intact glycoforms of various O-glycosites recently found in the S-RBD because they are expected to play a role in viral function and viral binding with ACE2.^17,22,23,33^ Therefore, new methods capable of the comprehensive structural elucidation of O-glycans are essential for understanding the functional roles of O-glycans on SARS-CoV-2 pathology and provide new avenues for rational therapeutic development.^28,34-36^

Conventional methods to analyze protein glycosylation rely on the bottom-up MS approach of analyzing glycopeptides obtained through enzymatic digestion.^31,37,38^ Although the bottom-up MS approach is a high throughput method for global proteome identification of glycosites and N-glycan heterogeneity from complex samples, the relative abundance of various intact glycoforms and the overall post-translational modification (PTM) compositions of different co-appearing ‘proteoforms’^39^ are lost.^40-42^ In contrast, by combining glycoproteomics with the top-down MS approach, which preserves the intact glycoprotein enabling high-resolution proteoform-resolved analysis,^43-45^ we could achieve the simultaneous characterization of the molecular structures, the site specificity, and the relative abundance of various glycoforms. Furthermore, by integrating native MS, which has recently emerged as a powerful structural biology tool to study protein structure-function relationships,^46-52^ with trapped ion mobility spectrometry (TIMS),^53-55^ we can investigate the gas phase structural variants to achieve the direct quantification of individual glycoproteoforms.

Here we developed a method for the molecularly detailed analysis of intact O-glycan proteoforms of the S-RBD by top-down MS (**Figure 1**). For the first time, we harness the capabilities of a hybrid TIMS quadrupole time-of-flight (QTOF) mass spectrometer (timsTOF Pro) (Figure 1C), which provides high resolving power and sensitivity for both selective and comprehensive ion mobility separations of various protein structural variants,^56-59^ and the ultrahigh-resolution of a 12T Fourier Transform Ion Cyclotron Resonance (FTICR) MS (Figure 1D), to comprehensively characterize the O-glycoforms of the S-RBD, including the exact glycan structures and relative molecular abundance (Figure 1E). We demonstrate that this native top-down MS approach can provide a high-resolution and wholistic proteoform-resolved landscape of diverse O-glycoforms to enable future structure-function studies of the S-RBD.

**Figure 1.**
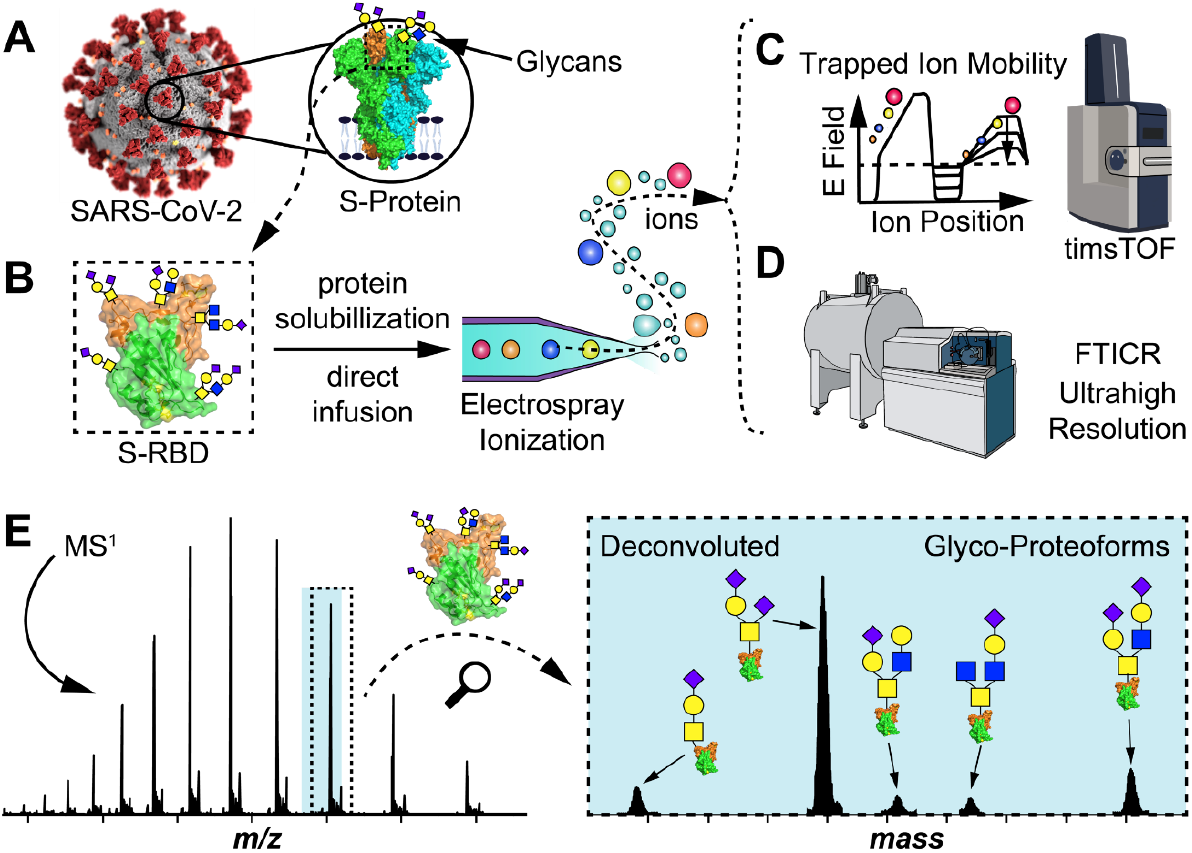
High-resolution glycoform characterization of intact S-RBD by top-down MS. (A-D) Illustration of the top-down glycoproteomics workflow for the comprehensive analysis of the S-RBD glycoforms. (A) The SARS-CoV-2 coronavirus features a surface S protein that possesses a glycosylated RBD (highlighted in the dashed box in B). (B) Intact glycoprotein analysis proceeds by directly infusing solubilized S-RBD and electrospraying S-RBD protein ions into either (C) a hybrid trapped ion mobility spectrometry (TIMS) MS device (timsTOF) for ion mobility MS analysis, or (D) an ultrahigh-resolution Fourier transform ion cyclotron (FTICR) MS. (E) Specific isolation of S-RBD proteoforms by top-down MS analysis, illustrating raw MS^1^ and corresponding deconvoluted protein spectrum, for structural characterizations of glycoforms. PDB: 6M0J.

## Results and Discussion

### Native TIMS-MS analysis of the S-Protein RBD

We first prepared a native S-RBD protein expressed from HEK 293 cells to perform native top-down MS for comprehensive glycoform analysis (**Figure 1**). After ensuring reproducible native S-RBD protein sample preparation (**Figure S1**), we used the TIMS, the front-end of the timsTOF Pro (see Methods for details), to achieve high resolution ion mobility separation of protein conformers. TIMS, similar to other quantitative ion mobility techniques, reports the rotationally average collision cross section (CCS) of proteins, which relates to their overall size and shape, and consequently can be used to evaluate changes in three-dimensional structure.^57^ However, ion mobility mapping (**Figures 2A**) and MS^1^ analysis (**Figure 2B**) reveal enormous complexity in the native S-RBD sample. The diffuse spread of the ion mobility spectrum suggests multiple co-existing S-RBD proteoforms potentially arising from multiple glycans (**Figure 2A**). Closer inspection of the ion mobility and native MS spectra reveals significant glycan heterogeneity likely due to the multiple N-and O-glycans on the native S-RBD (**Figures 2C,D**). The native S-RBD could not be resolved due to overlapping ion signals from all diverse glycoforms/proteoforms carrying different charges, similarly as observed in complex glycoproteins previously.^40,53^

**Figure 2.**
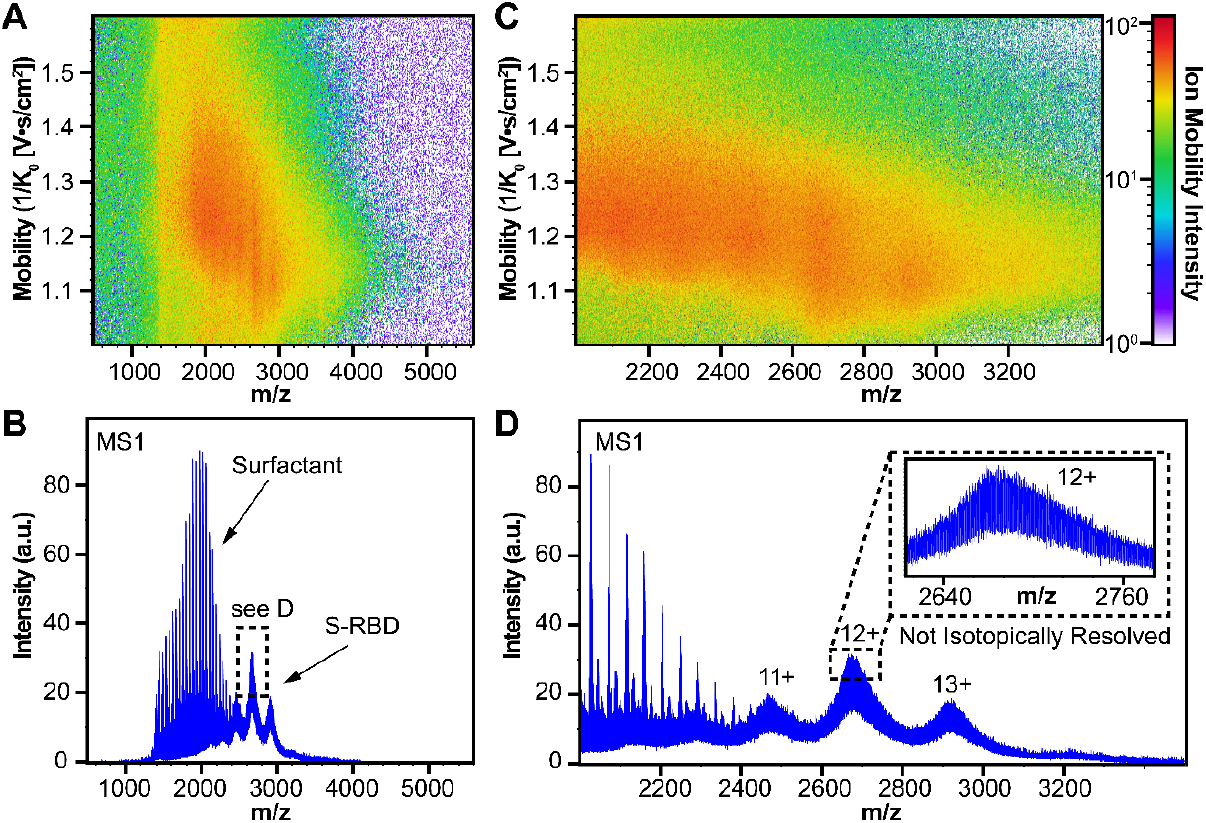
Native top-down MS of intact S-RBD expressed from HEK 293 cell line using TIMS-MS. (A-B) Ion mobility heat map (A) and native MS^1^ analysis (B) of the S-RBD. (C) Magnified ion mobility heat map for the S-RBD region from (A). The diffuse ion mobility spectrum suggests a diverse spread of multiple protein ions with various collisional cross sections crowded in the native spectrum. (D) Magnified MS^1^ from (B) with charge states of the native S-RBD annotated. The inset reveals that the large cluster of various proteoforms of the S-RBD inhibits its isotopic resolution.

The S protein is known to carry many complex-type N-glycans, with two glycosites (Asp331 and Asp343) previously reported and extensively characterized on the S-RBD.^21,22^ By contrast, the O-glycans of the S-RBD are less studied and their glycoforms poorly understood, despite the potential insights they may provide to understand viral structure-function relationship.^17,21-23^ Since our focus is on O-glycans of the S-RBD, we completely removed the N-glycans from the S-RBD using a native PNGase F treatment (see Methods for details) to minimize the interference posed by the enormous N-glycan heterogeneity (**Figure 3A**). The removal of N-glycans constitutes >10 kDa of molecular weight loss compared to the fully glycosylated S-RBD, as expected from the multiple proteoforms detected by TIMS analysis (**Figures 2A,C**). Fortunately, we found that after complete removal of N-glycans, the remaining O-glycans on the S-RBD were now fully resolvable by native MS (**Figure 3B**). Although the remaining PNGase F was found coexisting with the S-RBD, further optimization of the deglycosylation treatment yielded highly resolved charge states of S-RBD (**Figure 3C**). Deconvoluted MS analysis of the isotopically resolved charge states revealed multiple O-glycoforms with molecular weights ranging from 25.5 kDa to 26.8 kDa (**Figure 3C** inset).

**Figure 3.**
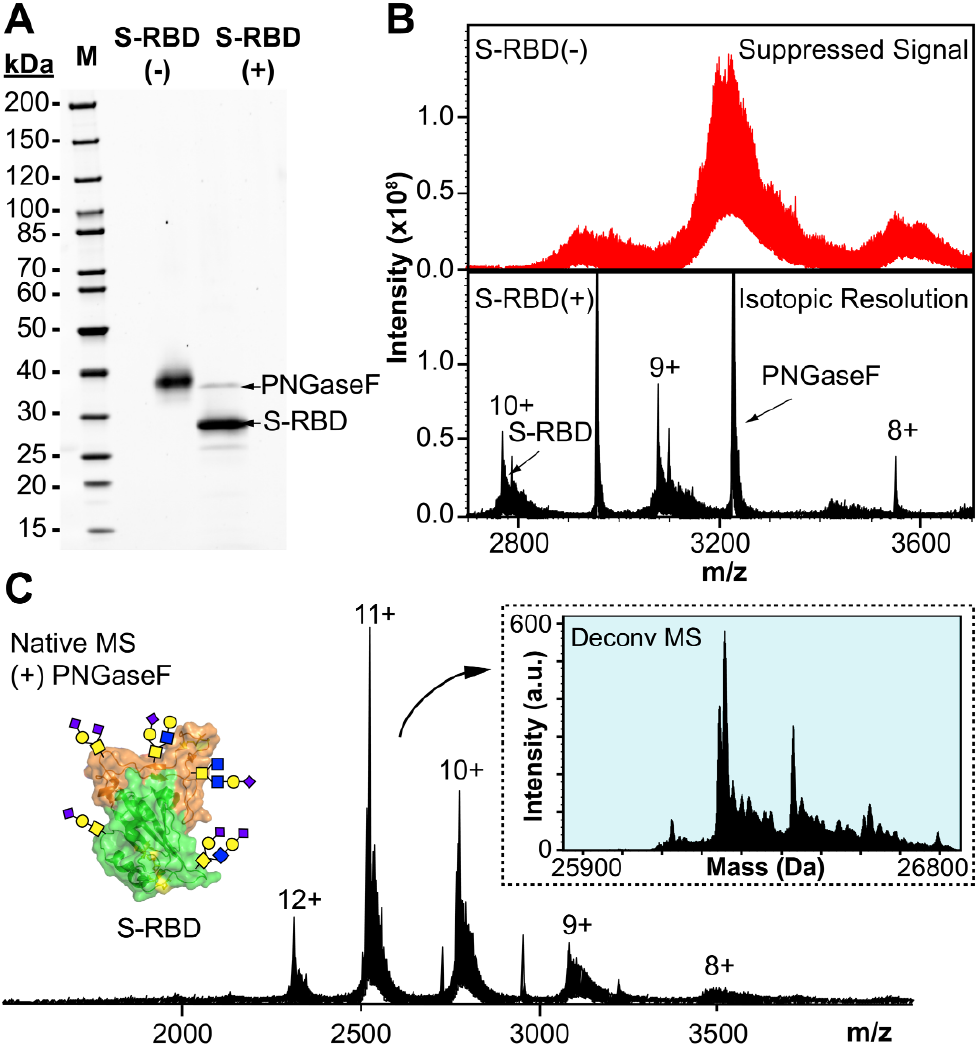
Isotopically resolved native top-down MS of intact S-RBD after N-glycan removal. (A) SDS-PAGE of S-RBD before (-) and after (+) PNGase F treatment. The gel lanes were equally loaded (200 ng) and staining was visualized by SYPRO Ruby. (B) Native MS of the S-RBD before (-) and after (+) PNGase F treatment showing improved ion signal and resolution, corresponding to the data shown in (A). (C) Isotopically resolved S-RBD obtained after optimization of PNGase F treatment. The protein charge states are labeled, and the inset shows the corresponding deconvoluted mass spectrum.

We next used TIMS to separate and analyze the various S-RBD glycoforms after the N-glycan removal by taking full advantage of the TIMS front-end of the timsTOF Pro instrument. After careful TIMS tuning and optimization of the ion mobility parameters (**Figure S2**), the timsTOF was sufficiently sensitive to allow high resolution ion mobility separation of intact S-RBD O-glycoforms (**Figure 4A, Figure S3**), in contrast to the poorly resolved native S-RBD (**Figure 2**). The TIMS analysis revealed two distinct protein gas-phase conformers separated in regional mobility between 1.1 to 1.45 1/K_0_. By interrogating each of the resolved mobility regions, MS^1^ analysis revealed that the two gas-phase conformers show shifts in the relative abundance of the S-RBD charge states (**Figure 4B**). The shift in protein charging along with the regional mobility shift together imply possible structural changes which can influence the protein ionization between the mobility regions. This difference in protein conformers was further investigated by plotting the corresponding ion mobility spectrum of the various regions against their calculated CCS values (**Figure 4C**).

**Figure 4.**
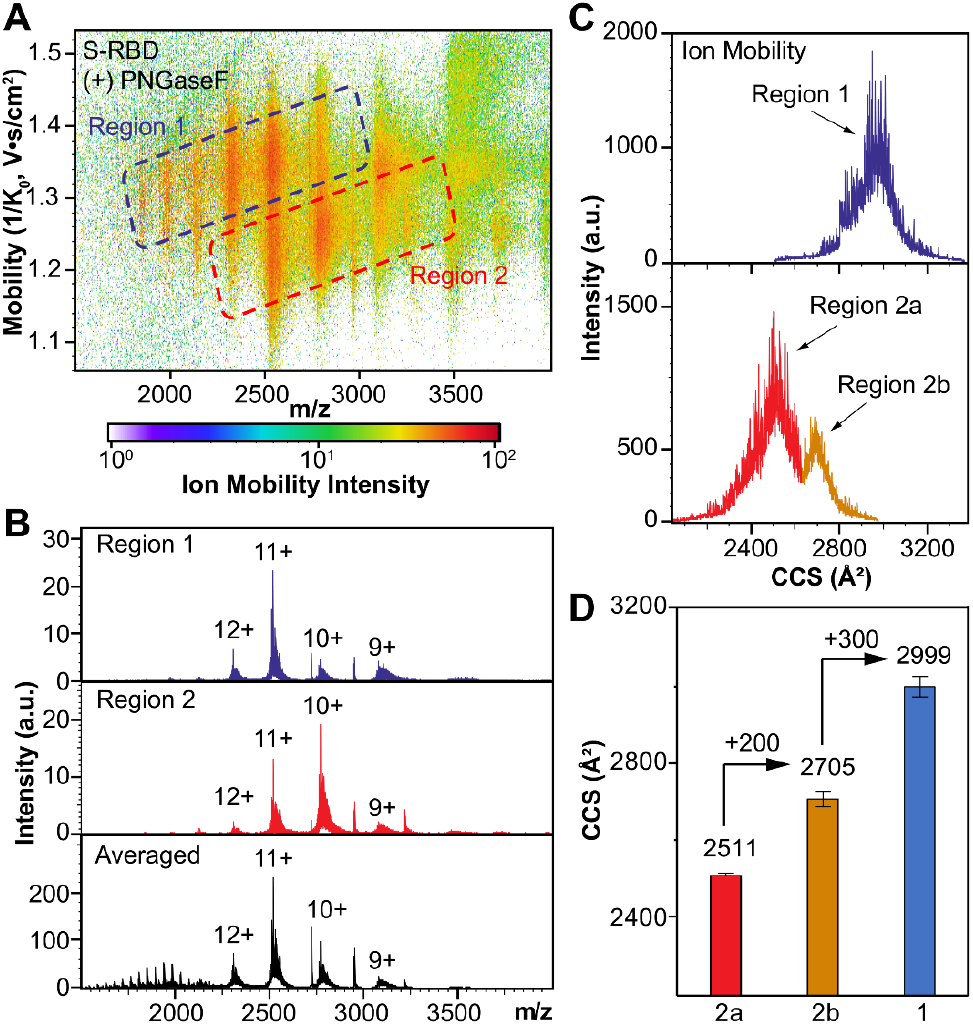
Ion mobility separation of intact S-RBD O-glycoforms by native TIMS-MS. (A) Ion mobility heat map of S-RBD after PNGase F treatment reveals distinct gas-phase conformers labeled as Region 1 (blue) and Region 2 (red). (B) Native MS^1^ analysis of the various ion mobility regions corresponding to (A). The MS^1^ spectrum averaged across the two mobility regions is shown for reference. (C) Ion mobility spectrum corresponding to the data shown in (A). The two distinct peaks in Region 2 are illustrated as Region 2a (red) and Region 2b (orange). (D) Collision cross section (CCS) of the most abundant S-RBD ion (11+ charge state) ion mobility regions corresponding to (C). CCS data (D) are representative of *n* = 3 independent experiments with error bars indicating the standard deviation of the mean.

Applying this method, we further found two distinct S-RBD O-glycoform mobility regions in the mobility Region 2 (**Figure 4C** lower panel). The smallest mobility region of the S-RBD O-glycoforms (Region 2a, 2511±5 Å^2^) is slightly higher than the theoretical value (∼2200 Å^2^) calculated using the IMPACT method^60^ from the S-RBD X-ray crystal structure without glycosylation.^61^ This discrepancy is likely a result of the influence of the additional O-glycans.^62^ CCS measurements of the various ion mobility regions of the S-RBD revealed progressively larger structures between the highlighted mobility regions and further illustrates the separation of the distinct gas-phase conformers (**Figure 4D**). Region 1 (2999±26 Å^2^) shows the largest calculated CCS value compared to Region 2b (2705±25 Å^2^) and Region 2a (2511±5 Å^2^), implying that the S-RBD glycoforms in Region 1 experience additional protein unfolding which increases the overall protein conformer size and signal abundance under electrospray ionization.^63^ The variations in S-RBD O-glycoforms CCSs are mobility region specific and consistent with protein conformer changes, illustrating the potential of this approach for investigating the gas-phase structural variations of glycoproteins.

### Comprehensive characterization of S-RBD O-glycoforms by high resolution top-down MS

Although S-RBD glycoforms can be inferred from the timsTOF analysis, we cannot assign the glycan structures or occupancy from only the results presented in **Figures 3,4** due to the mass degeneracy and microheterogeneity of O-glycans.^40,62^ To achieve in-depth glycoform analysis, we further utilized an ultrahigh-resolution 12T FTICR capable of baseline and isotopically resolving the S-RBD O-glycoforms for MS/MS analysis (see Methods for details, **Figure S4**). Following further MS^1^ analysis, individual glycoforms were directly visualized with high resolution (**Figure 5A,B**). Following specific quadrupole isolation centered at appropriate *m/z* widths to capture individual S-RBD proteoforms (**Figure 5C**), the resolved intact glycoforms can be characterized by MS/MS using a combination of fragmentation strategies to achieve confident protein sequence assignments (**Figure 5D**).

**Figure 5.**
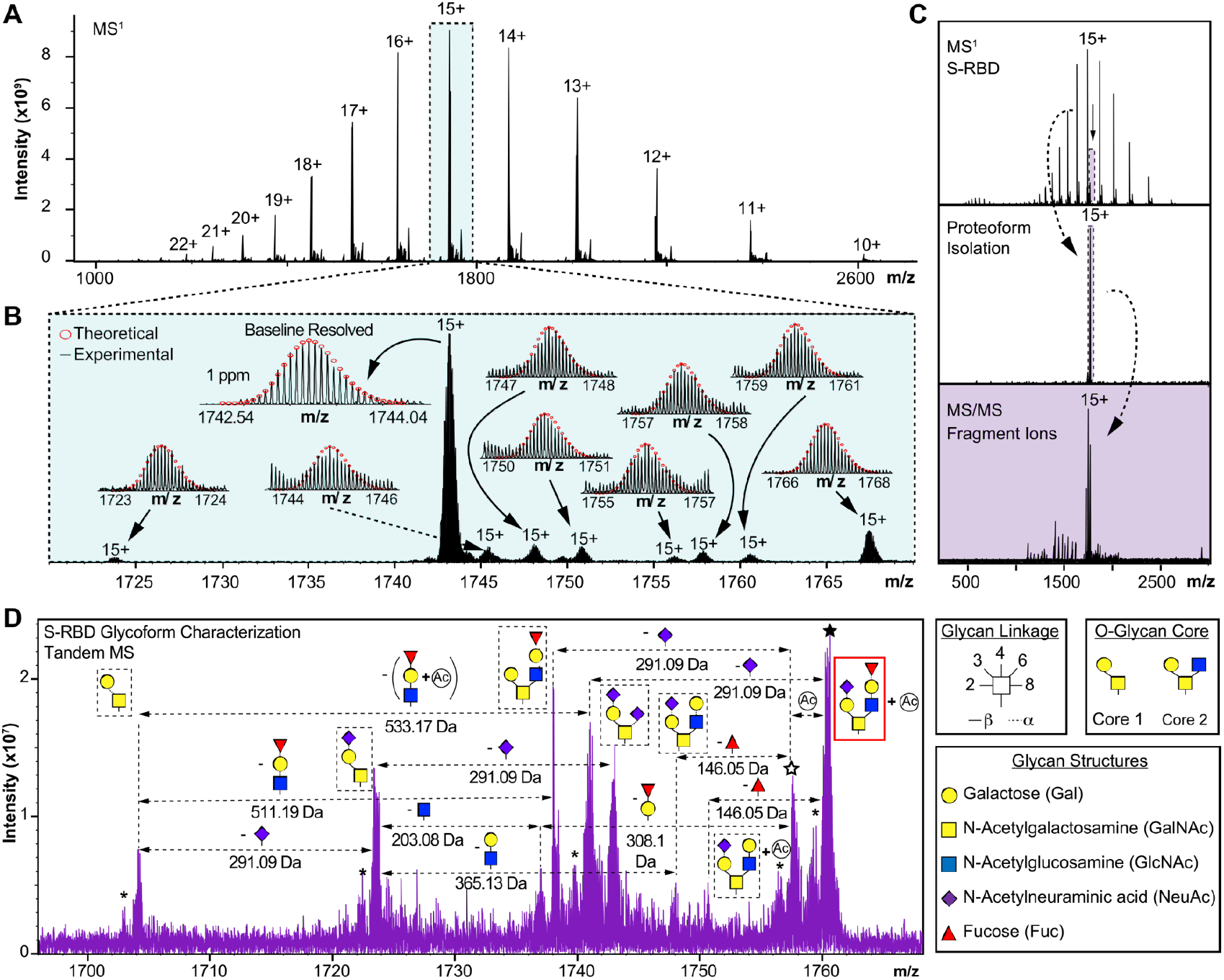
S-RBD O-glycoform proteoform analysis by high resolution top-down MS. (A) MS^1^ analysis of S-RBD O-glycoforms following denaturation. (B) Magnified MS^1^ from (A) showing isotopically and baseline resolved S-RBD proteoforms allowing for accurate determination of individual intact glycoforms. The most abundant charge state (15+) is isolated. Theoretical isotope distributions (red circles) are overlaid on the experimentally obtained mass spectrum to illustrate the high mass accuracy. All individual ion assignments are within 1 ppm from the theoretical mass. (C) Illustration of the isolation of a specific S-RBD glycoform using ultrahigh resolution FTICR to enable glycan site and structure characterization from intact glycoforms. (D) MS/MS characterization of S-RBD proteoform isolated from quadrupole window centered at 1760.5 *m/z*. The assignments of glycan structures are marked in the spectrum with the legends shown on the right side. Glycoform characterization reveals the specific S-RBD proteoform to have Core 2 type GalNAcGal(GalNeuAc)(GlcNAcGalFuc) glycan. Neutral loss glycan products are labeled. N-terminal acetylation (Ac) is labeled and corresponds to a +42 Da mass shift. The solid star represents the 15+ charge state precursor ion corresponding to 1760.5 *m/z*, and the hollow star represents the 15+ charge state precursor ion corresponding to 1757.7 *m/z* (-Ac). The asterisk “*” denotes an oxonium ion loss.

By applying this method on all detectable S-RBD proteoforms, a suite of MS/MS spectra were generated for sequence analysis (**Figure S5**). MS/MS data were output from the Bruker DataAnalysis software and analyzed in targeted protein analysis mode using MASH Explorer.^64^ This approach allowed for the direct characterization of the glycan structures and their microheterogeneity to reveal multiple S-RBD glycoforms with Core 1 and Core 2 O-glycan structures (**Figure 5**, and full details in **Figures S6-11**). This high-resolution method enables the structural characterization of glycoforms that have not been previously observed. As shown for one such example in **Figure 5D** (highlighted in the red box on the right), neutral loss mapping revealed a Core 2 (GlcNAcβ1-6(Galβ1-3)GalNAc-Ser/Thr) glycoform (15+ most abundant charge state, centered at 1760.5 *m/z*) with a GalNAcGal(GalNeuAc)(GlcNAcGalFuc) structure at Thr323. The site assignment of this glycan to Thr323 agrees with previous studies reporting putative assignments of other S protein O-glycans at the same site,^22^ which is reasonable given the presence of an adjacent proline at residue 322, increasing the likelihood of an O-glycosylation site. Direct visualization of MS/MS fragment ions obtained from collision-induced dissociation (CID) along with the corresponding proteoform sequence mapping confirms the glycan structure assignment with high mass accuracy (**Figure S6**). Additionally, the GalNAcGal(GalNeuAc)(GlcNAcGalFuc) related proteoform without N-terminal acetylation (15+ most abundant charge state, centered at 1757.7 *m/z*) was also characterized (**Figure S6**). Therefore, the high resolution glycoform mapping presented in **Figure 5** illustrates a unique strength of this native top-down MS approach to achieve isotopically resolved and high accuracy characterization of highly heterogenous glycoproteins, a major challenge in intact glycoprotein analysis.^40,65^

We also identified and characterized core 1 (Galβ1-3GalNAc-Ser/Thr) O-glycan structures such as GalNAcGalNeuAc (15+ most abundant charge state, centered at 1723.8 *m/z*) (**Figure S7)**, and GalNAcGal(NeuAc)_2_ (15+ most abundant charge state, centered at 1743.2 *m/z*) (**Figure S8**). Interestingly, these two Core 1 O-glycans were previously reported for the S-RBD as potential modifications, however the previous studies were not able to resolve the exact glycoforms due to the challenges arising from inferring intact glycoprotein structures from peptide digests and the signal low abundance of O-glycans under conventional MS analysis.^22,23,37^ However, our native top-down MS strategy enables us to unambiguously reveal the exact glycoform corresponding to each of these Core 1 O-glycans. Two additional Core 2 glycan structures were identified GalNAcGal(GalNeuAc)(GlcNAcGal) (15+ most abundant charge state, centered at 1748.1 *m/z*, with additional N-terminal acetylated proteoform found at 1750.5 *m/z*) (**Figures S9**,**10**) and GalNAcGal(GalNeuAc)(GlcNAcGalNeuAc) (centered at 1767.6 *m/z*) (**Figure S11**).

This native top-down MS approach enables the proteoform-resolved characterization of S-RBD O-glycoforms to identify and characterize multiple S-RBD O-glycoforms as well as a novel fucose-containing glycoform with Core 2 GalNAcGal(GalNeuAc)(GlcNAcGalFuc) glycan structure (**Figure 6A**). **Figure 6A** shows all of the 7 identified O-glycoforms of the S-RBD, which are also listed in the table in **Figure 6B**. As a unique advantage of the top-down MS approach, the molecular abundance of each intact protein glycoform can be relatively quantified. We found that the relative abundance of Core 1 to Core 2 S-RBD O-glycan structures was roughly 70:30, with the Core 1 GalNAcGal(NeuAc)_2_ being the most abundant O-glycoform (∼67% relative abundance) (**Figure 6B**). This ability to unambiguously elucidate the structure of a specific S-RBD glycoform with high accuracy and quantify its relative abundance demonstrates the distinct advantages of this native top-down MS strategy over glycopeptide-based bottom-up MS approaches.^47,62^ Together, this proteoform-resolved intact glycoprotein analysis strategy enables the simultaneous characterization of O-glycan structures of the S-RBD and its microheterogeneity, including structure and relative molecular abundance (**Figure 6C**). Importantly, the accurate determination of the relative abundance of intact glycoforms provides the technical foundation to understand the functional significance of these distinct glycoforms of S protein in the future. Although these identified O-glycoforms are specifically for S-RBD expressed from HEK293 systems, the HEK293 cell expression system is known to reflect the glycosylation sites expected for the viron^21- 23,27^ and is currently the antigen of choice for vaccine candidates and virus functional studies.^66,67^

**Figure 6.**
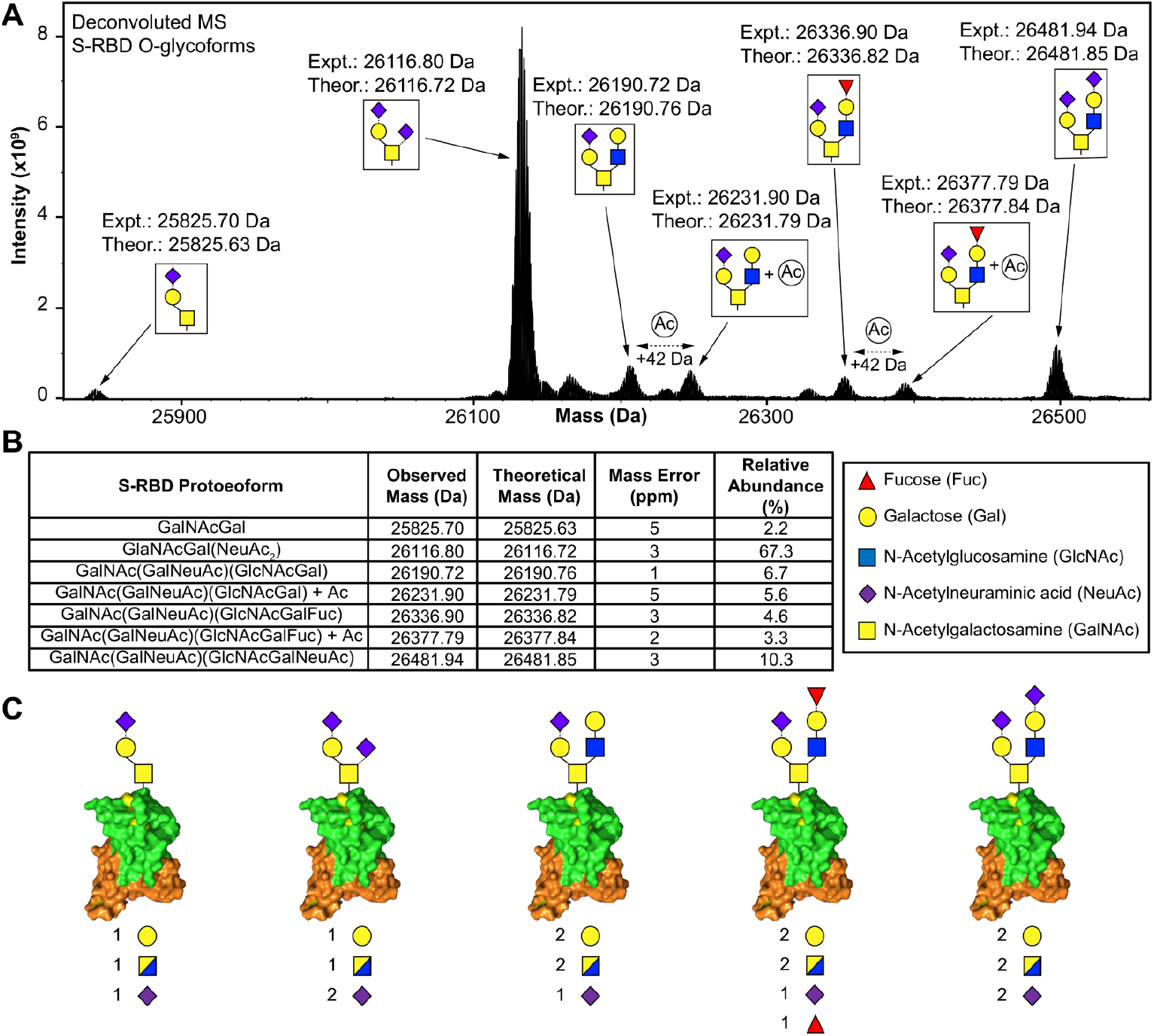
Complete characterization of the structures and relative abundance of S-RBD O-glycoforms by native top-down MS. (A) Deconvoluted MS of the S-RBD O-glycoforms showing all seven intact glycan assignments revealed by top-down MS/MS. Experimental and theoretical mass assignments are illustrated for each glycoform. (B) Summary of the O-glycoform characterization for the S-RBD with the relative abundance and mass accuracy of each identified glycoform. Masses are reported as the monoisotopic mass. (C) Illustration of S-RBD O-glycan microheterogeneity identified by specific proteoform isolation and sequencing.

## Conclusion

In summary, the combination of native trapped ion mobility MS from timsTOF and native top-down MS from ultrahigh-resolution FTICR enables the deep and comprehensive analysis of the intact glycoforms of the S-RBD, for the first time. Importantly, we report the complete structural characterization of seven S-RBD glycoforms, along with a Core 2 fucosylated glycan structure not previously reported. Moreover, the relative molecular abundances of each of these intact glycoforms can be quantified, providing new insights into the complex landscape of S-RBD O-glycosylation. Our proteoform-resolved method provides complete structural characterization of O-glycan microheterogeneity and complex glycoproteins. This native top-down mass spectrometry approach not only enables the comprehensive analysis of the S-RBD and the molecular signatures of emerging S-RBD variants of SARS-CoV-2 virus toward the structure-function studies in the future, but also can help catalyze the studies of other O-glycoproteins in general.

## Materials and Methods

### Materials and Reagents

All chemicals and reagents were purchased from MilliporeSigma (Burlington, MA, USA) and used as received without further purification unless otherwise noted. Aqueous solutions were made in nanopure deionized water (H_2_O) from Milli-Q® water (MilliporeSigma). Ammonium acetate (AA) was purchased from Fisher Scientific (Fair Lawn, NJ). Bradford protein assay reagent was purchased from Bio-Rad (Hercules, CA, USA). 10% gel (10 or 15 comb well, 10.0 cm × 10.0 cm, 1.0 mm thick) for SDS-Polyacrylamide Gel Electrophoresis (SDS-PAGE) was home-made. Thermo Scientific™ Cimarec+™ stirring hotplate were purchased from ThermoFisher Scientific. Amicon, 0.5 mL cellulose ultra centrifugal filters with a molecular-weight cutoff (MWCO) of 10 kDa were purchased from MilliporeSigma. The recombinant SARS-CoV-2 spike regional-binding domain protein expressed in the HEK 293 cell line was purchased from Sino Biological Inc. (cat. 40592-VNAH).

### Complete N-glycan removal for O-glycan protein analysis

Peptide-N-glycosidase F (PNGase F) (New England Biolabs Inc., Cat. P0704S) was used for complete removal of N-linked oligosaccharides. Briefly, 20 μg of protein sample was buffer exchanging into 150 mM AA (pH = 7.4) solution by washing the sample five times through a 10 kDa Amicon ultra centrifugal filters (MilliporeSigma, Burlington, MA, USA). PNGase F (5 U) was then added to the protein solution and samples were incubated at 37 °C for 12 h on a Thermo Scientific™ Cimarec+™ stirring hotplate.

### Sample preparation

Native protein samples were prepared by buffer exchanging into 150 mM AA solution by washing the sample five times through a 10 kDa Amicon ultra centrifugal filters (MilliporeSigma, Burlington, MA, USA). The protein sample was then diluted to 20 μM in 150 mM AA prior to native top-down MS analysis. Denatured protein samples were reduced using 20 mM tris(2-carboxyethyl)phosphine hydrochloride (TCEP), and followed by buffer exchanging into 0.1 % formic acid solution by washing the sample five times through a 10 kDa Amicon ultra centrifugal filters (MilliporeSigma, Burlington, MA, USA). The protein sample was then diluted to 20 μM in 50:50:0.1 (acetonitrile: water: formic acid) prior to top-down MS analysis. Protein samples subjected to N-glycan removal were prepared similarly following PNGase F reaction.

### Top-down MS analysis

Intact protein samples were analyzed by nano-electrospray ionization via direct infusion using a TriVersa Nanomate system (Advion BioSciences, Ithaca, NY, USA) coupled to a solariX XR 12-Tesla Fourier Transform Ion Cyclotron Resonance mass spectrometer (FTICR-MS, Bruker Daltonics, Billerica, MA). For the nano-electrospray ionization source using a TriVersa Nanomate, the desolvating gas pressure was set at 0.45 PSI and the voltage was set to 1.2 to 1.6 kV versus the inlet of the mass spectrometer. Source dry gas flow rate was set to 4 L/min at 124 °C. For the source optics, the capillary exit, deflector plate, funnel 1, skimmer voltage, funnel RF amplitude, octopole frequency, octopole RF amplitude, collision cell RF frequency, and collision cell RF amplitude were optimized at 190 V, 200 V, 100 V, 35 V, 250 Vpp, 2 MHz, 490 Vpp, 2MHz, and 2000 Vpp, respectively. Mass spectra were acquired with an acquisition size of 4M, in the mass range between 200-4000 *m/z* (with a resolution of 530,000 at 400 *m*/*z*), and a minimum of 250 scans were accumulated for each sample. Ions were accumulated in the collision cell for 0.100 s, and a time of flight of 1.500 ms was used prior to their transfer to the ICR cell. For collision-induced dissociation (CID) tandem MS (MS/MS) experiments, the collision energy was varied from 10 to 30V, ion accumulation was optimized at 400 ms, and file size was 4M. Tandem mass spectra were output from the DataAnalysis 4.3 (Bruker Daltonics) software and analyzed using MASH Explorer.^64^

### Native top-down MS analysis

A timsTOF Pro mass spectrometer (Bruker Daltonics, Bremen, Germany) was couple to a Bruker nanoElute LC system (Bruker Daltonics, Bremen, Germany). Samples were directly infused using the nanoElute, injecting 5 μL of native protein sample with a flow rate of 1 μL/min. For the MS inlet, the endplate offset and capillary voltage were set to 500 V and 3800 V, respectively. The nebulizer gas pressure (N_2_) was set to 1.5 bar, with a dry gas flow rate of 6 L/min at 180 °C. The tunnel out, tunnel in, and TOF vacuum pressures were set to 0.8584 mBar, 2.577 mBar, and 1.752E-07 mBar. To calibrate the MS and trapped ion mobility spectrometry (TIMS) device, Agilent tune mix was directly infused to provide species of known mass and reduced mobility.^37^ For MS calibration, the MS resolution for the most abundant calibrant signal, 1222 *m/*z, was 62,000. Calibrant points at 922, 1222, and 1522 *m/z* were used for TIMS calibration. The TIMS resolution for the most abundant calibrant signal, 1222 *m/*z, was 80.2 CCS/ΔCCS. IMS tunnel voltage deltas were optimized at -20 V, -120 V, 70 V, 200 V, 0 V and 100 V for Δ1 to Δ6, respectively. TIMS funnel 1 RF was set to 350 Vpp, and TIMS collision cell energy was set to 250 V. An IMS imeX accumulation time and cycle ramp time of 400 ms was found to yield optimal TIMS resolving power. To facilitate high MS signal intensity, the TIMS accumulation time was locked to a 100% duty cycle, or 400 ms accumulation time. In the MS transfer optics the funnel 1 RF, funnel 2 RF, deflection delta, in-source collision induced dissociation (isCID) energy, multipole RF, and quadrupole ion energy were optimized at, 350 Vpp, 500 Vpp, 55 V, 0 eV, 800 Vpp, and 4 eV, respectively. For MS1 spectral collection, the quadrupole low mass was set to 1000 *m/z* with a scan range of 200 to 5000 *m/z*. Collision energy was set to 6 eV, with 3500 Vpp collision cell RF, a 100 μs transfer time, and a pre pulse storage time of 35 μs. For MS2 spectral collection the quadrupole low mass was set to 200 *m/z* with a scan range of 200 to 5000 *m/z*. The collision cell RF was set to 3500 Vpp, a 100 μs transfer time, and a pre pulse storage time of 20 μs were used. Mass spectra were output from the DataAnalysis 5.3 (Bruker Daltonics) software and analyzed using MASH Explorer.^64^

### Data Analysis

All data were processed and analyzed using Compass DataAnalysis 4.3/5.3 and MASH Explorer.^64^ Maximum Entropy algorithm (Bruker Daltonics) was used to deconvolute all mass spectra with the resolution set to 80,000 for the timsTOF Pro or with instrument peak width set to 0.05 for the 12T FTICR. Sophisticated Numerical Annotation Procedure (SNAP) peak-picking algorithm (quality factor: 0.4; signal-to-noise ratio (S/N): 3.0; intensity threshold: 500) was applied to determine the monoisotopic mass of all detected ions. The relative abundance for each protein isoform was determined using DataAnalysis. To quantify protein modifications, the relative abundances of specific modifications were calculated as their corresponding percentages among all the detected protein forms in the deconvoluted averaged mass. MS/MS data were output from the DataAnalysis software and analyzed using MASH Explorer^64^ for proteoform identification and sequence mapping. All the program-processed data were manually validated. For peak picking, the sophisticated numerical annotation procedure (SNAP) algorithm from Bruker DataAnalysis 5.3 was used with a quality threshold of 0.5 and an S/N lower threshold of 3. All fragment ions were manually validated using MASH Explorer. Peak extraction was performed using a signal-to-noise ratio of 3 and a minimum fit of 60%, and all peaks were subjected to manual validation. All identifications were made with satisfactory numbers of assigned fragment (>10), and a 25-ppm mass tolerance was used to match the experimental fragment ions to the calculated fragment ions based on amino acid sequence. For ion mobility analysis by the timsTOF Pro, the collisional cross section (CCS) in Å^2^ for a species of interest was calculated via the Mason-Schamps equation (Equation 1):

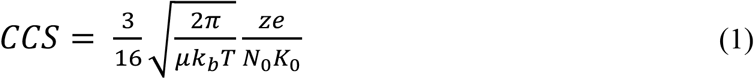

Where *μ* is the reduced mass of the ion-gas pair 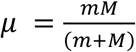, where m and M are the ion and gas particle masses), *k*_*b*_ is Boltzmann’s constant, *T* is the drift region temperature, *z* is the ionic charge, *e* is the charge of an electron, *N*_*0*_ is the buffer gas density, and *K*_*0*_ is the reduced mobility. Equation 1 was selected to agree with previously published CCS calculations.^57,58,68^ Theoretical CCS were calculated using the IMPACT method.^60^

## Supporting Information

The Supporting Information is available free of charge on the ACS Publications website.

## Acknowledgement

This research is supported by NIH R01 GM117058 (to S.J. and Y.G.). Y.G. would like to acknowledge NIH R01 GM125085, R01 HL096971, and S10 OD018475. We would like to thank Guillaume Tremintin, Yue Ju, Conor Mullins, Michael Greig, Gary Kruppa, Paul Speir and Rohan Thakur of Bruker Daltonics for their helpful discussion and provision of the Bruker timsTOF Pro used in this work.

