## Supplementary Materials for "Structural O-Glycoform Heterogeneity of the SARS-CoV-2 Spike Protein Receptor-Binding Domain Revealed by Native Top-Down Mass Spectrometry"

**Table of Contents** **Page**

Figures S1-11 2-13

### Supplementary Figures

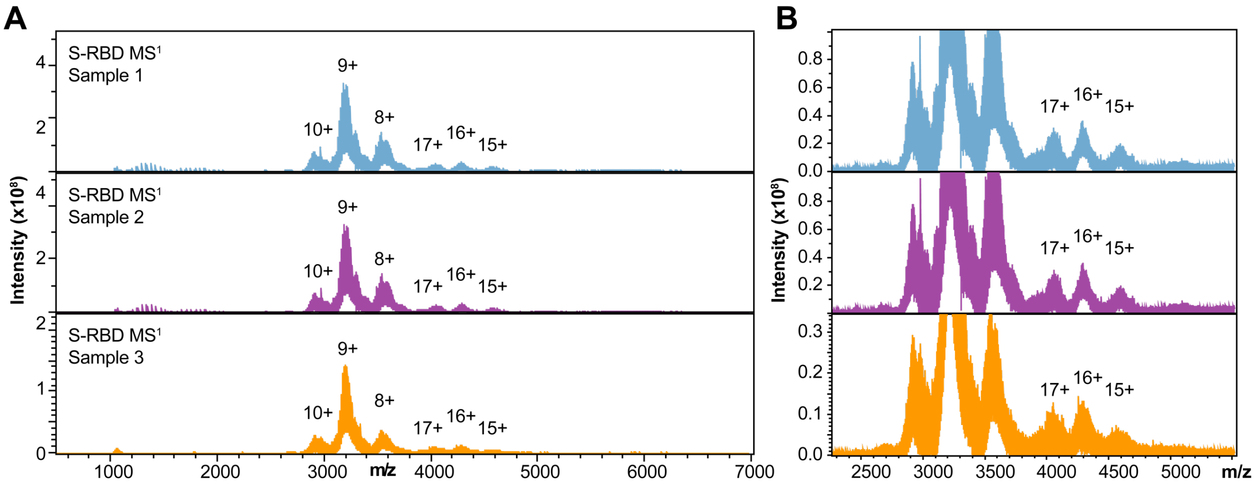


**Figure S1. Reproducibility of S-RBD native MS analysis.** (A) $MS^1$ of the native S-RBD sample. Three samples corresponding to different sample preparations are shown to demonstrate the reproducibility. (B) Close inspection of the $MS^1$ reveals potential dimer of S-RBD resolved by native MS. Isotopic signal is suppressed by the large glycan heterogeneity.

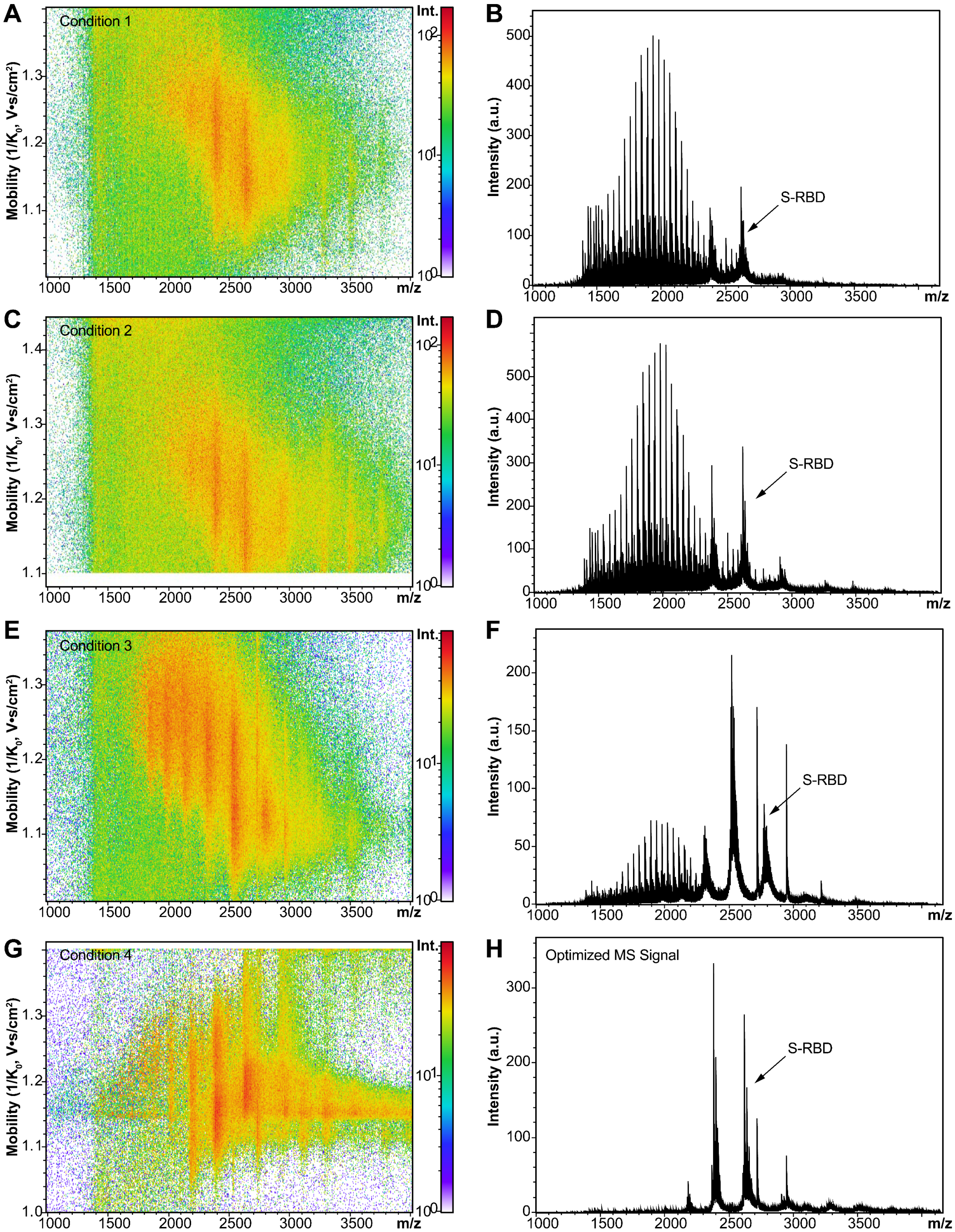


**Figure S2. Optimization of trapped ion mobility (TIMS) parameters for high resolution native S-RBD collision cross section (CCS) analysis.** (A,C,E,G) Ion mobility heat map showing distribution of $1/K_0$ (protein ion mobility) as a function of *m/z* for MS[1] of the native S-RBD sample after PNGase F treatment. Each of the (A), (C), (E), and (G) correspond to different TIMS parameter tuning. (B, D, F, G) MS[1] corresponding to the ion mobility heat maps shown in (A), (C), (E), and (G), respectively. Condition 4 with a collision energy of 10 eV was found to provide the best tradeoff between ion mobility resolution and MS[1] resolution, while preserving the native protein sample.

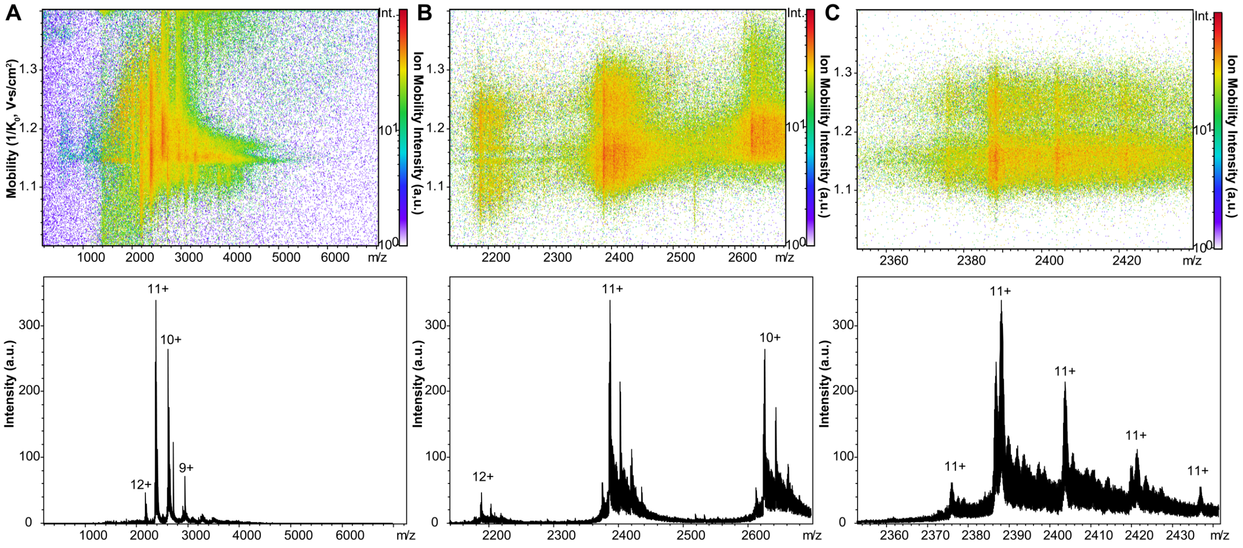


**Figure S3. High resolution ion mobility analysis of native S-RBD.** (A) Ion mobility heat map and corresponding native MS[1] of S-RBD after PNGase F treatment following optimized TIMS parameters from Figure S2. (B) Zoom-in of the various protein charge states (12+, 11+, and 10+) showing corresponding ion mobility heat map. (C) Focusing on a single charge state (11+), the isotopically resolved MS signal reveals various S-RBD proteoforms reflected as distinct mobility values.

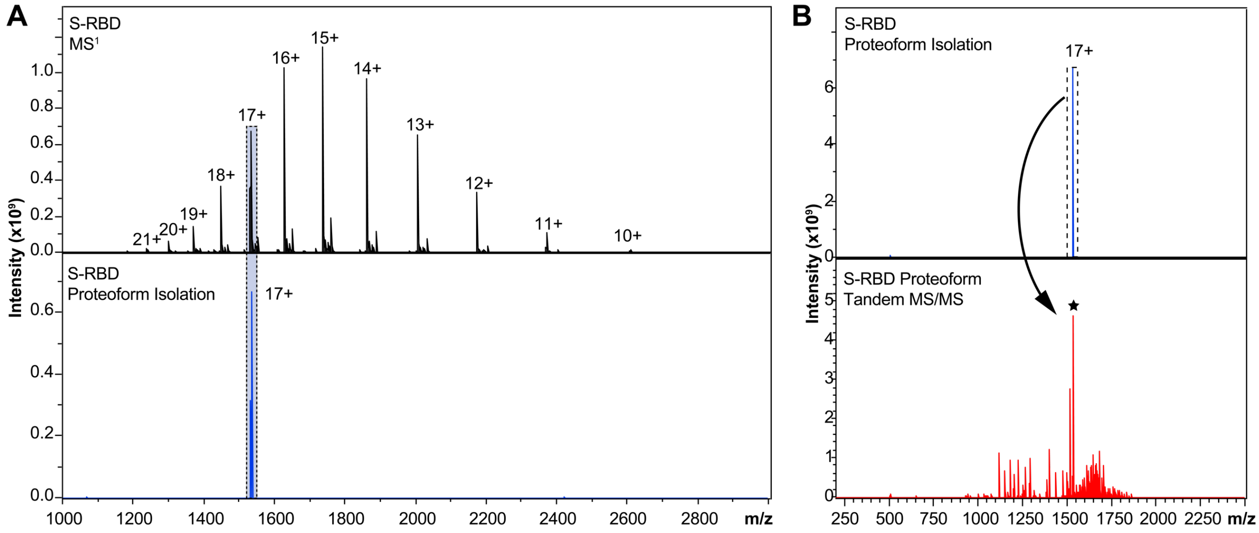


**Figure S4. Illustration of specific S-RBD proteoform characterization.** (A) $MS^1$ of S-RBD after PNGase F treatment showing baseline isotopic isolation of specific S-RBD proteoform at 17+ charge state. (B) Tandem MS/MS characterization of the isolated S-RBD proteoform corresponding to (A). The star represents the 17+ charge state precursor ion.

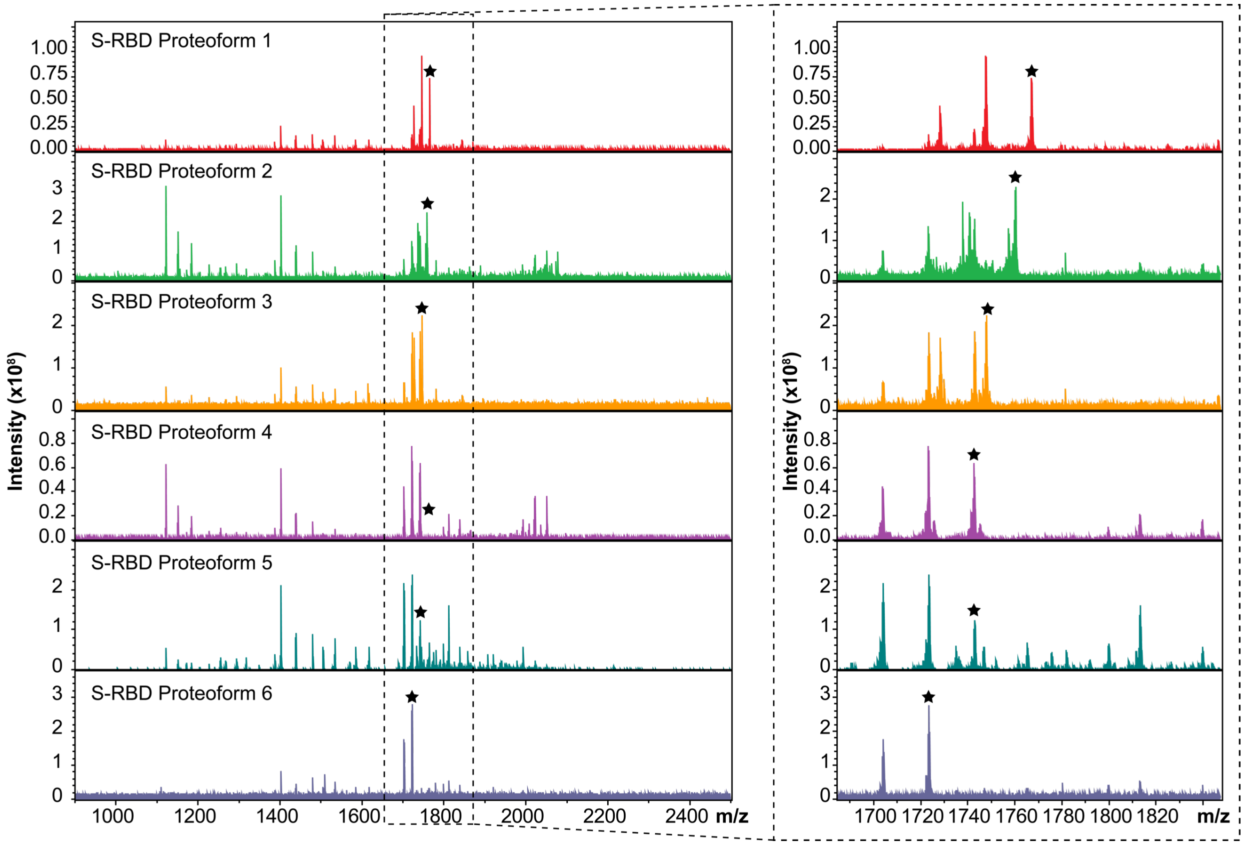


**Figure S5.** Tandem MS/MS suite illustrating the characterization of various S-RBD proteoforms after specific isolation. The MS/MS spectra of S-RBD proteoforms 1 to 7 are obtained from charge state 15+ of the S-RBD and after specific quadrupole isolation with windows centered at 1767.6 *m/z*, 1760.5 *m/z*, 1748.1 *m/z*, 1745 *m/z*, 1743.2 *m/z*, and 1723.8 *m/z*, respectively. Isolation widths were set to 2 *m/z*. The solid star represents the specific 15+ charge state precursor ion.

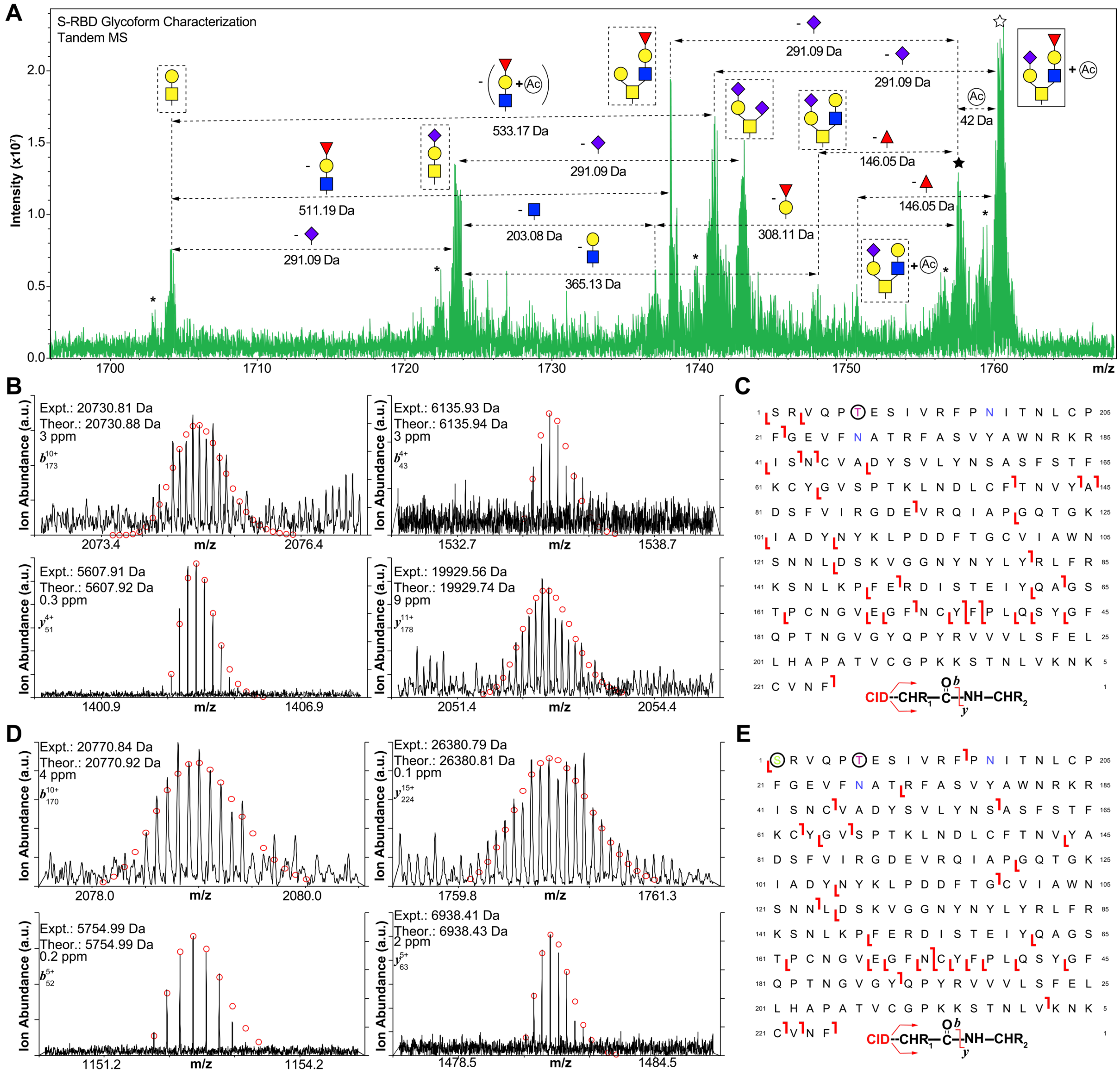


**Figure S6.** (A) MS/MS characterization of S-RBD proteoforms isolated from quadrupole window centered at 1757.7 *m/z* and 1760.5 *m/z*, corresponding to the MS[1] shown in Fig. 5. Glycoform characterization of the isolated 1760.5 *m/z* and 1757.7 *m/z* proteoforms reveal the specific S-RBD proteoform to have Core 2 type GalNAcGal(GalNeuAc)(GlcNAcGalFuc) glycan with and without a +42 Da mass shift corresponding to acetylation (Ac), respectively. Neutral loss glycan products are labeled. The solid star represents the 15+ charge state precursor ion corresponding to 1757.7 *m/z*, and the hollow star represents the 15+ charge state precursor ion corresponding to 1760.5 *m/z* (+Ac). The asterisk "*" denotes an oxonium ion loss. (B) Representative CID fragment ions ($b_{173}^{10+}$, $b_{43}^{4+}$, $y_{51}^{4+}$, and $y_{178}^{11+}$) obtained from S-RBD proteoform isolated from 1757.7 *m/z* precursor with glycosite at Thr323. (C) Fragmentation mapping of the specific S-RBD proteoform corresponding to (B). (D) Representative CID fragment ions ($b_{170}^{10+}$, $y_{224}^{15+}$, $b_{52}^{5+}$, and $y_{63}^{5+}$) obtained from S-RBD proteoform isolated from 1760.5 *m/z* precursor with glycosite at Thr323. Theoretical ion distributions are indicated by the red dots and mass accuracy errors are listed for each fragment ion. (E) Fragmentation mapping of the specific S-RBD proteoform corresponding to (D), showing N-terminal acetylation (Ac). Amino acid sequence (Arg319-Phe541) was based on the entry name P0DTC2 (SPIKE_SARS2) obtained from the UniProtKB sequence database. N-terminal Ser is due to removal of signal peptide tPA following cell expression. The blue N (Asn) denotes deamidation following PNGase F treatment.

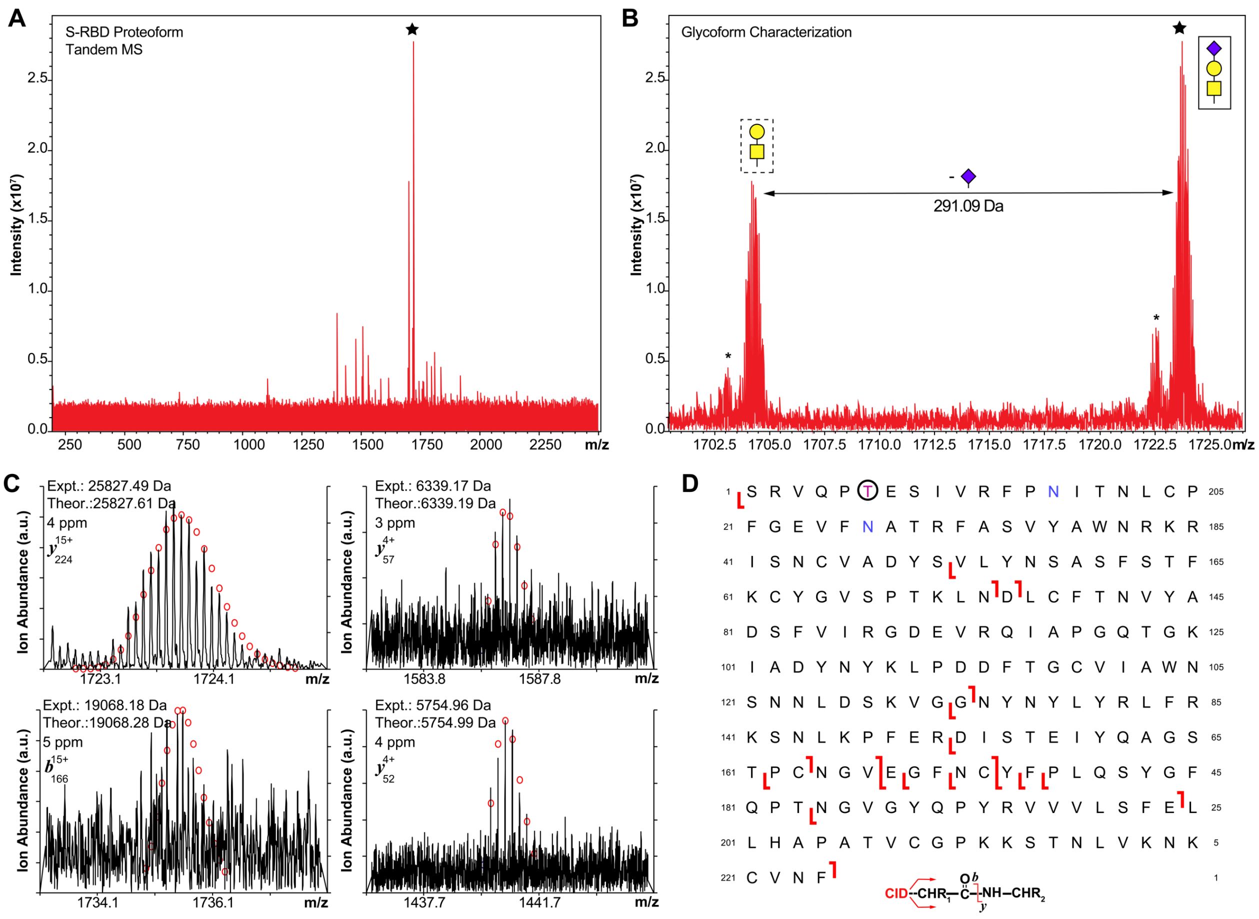


**Figure S7.** (A) MS/MS characterization of S-RBD proteoform isolated from quadrupole window centered at 1723.8 *m/z*, corresponding to the MS[1] shown in Fig. 5. (B) Glycoform characterization reveals the specific S-RBD proteoform to have Core 1 type GalNAcGalNeuAc glycan. Neutral loss glycan products are labeled. The star represents the 15+ charge state precursor ion and the asterisk "*" denotes an oxonium ion loss. (C) Representative CID fragment ions ($y_{224}{}^{15+}$, $y_{57}{}^{4+}$, $b_{166}{}^{15+}$, and $y_{52}{}^{4+}$) obtained from S-RBD with glycosite at Thr323. Theoretical ion distributions are indicated by the red dots and mass accuracy errors are listed for each fragment ion. (D) Fragmentation mapping of the specific S-RBD proteoform. Amino acid sequence (Arg319-Phe541) was based on the entry name P0DTC2 (SPIKE_SARS2) obtained from the UniProtKB sequence database. N-terminal Ser is due to removal of signal peptide tPA following cell expression. The blue N (Asn) denotes deamidation following PNGase F treatment.

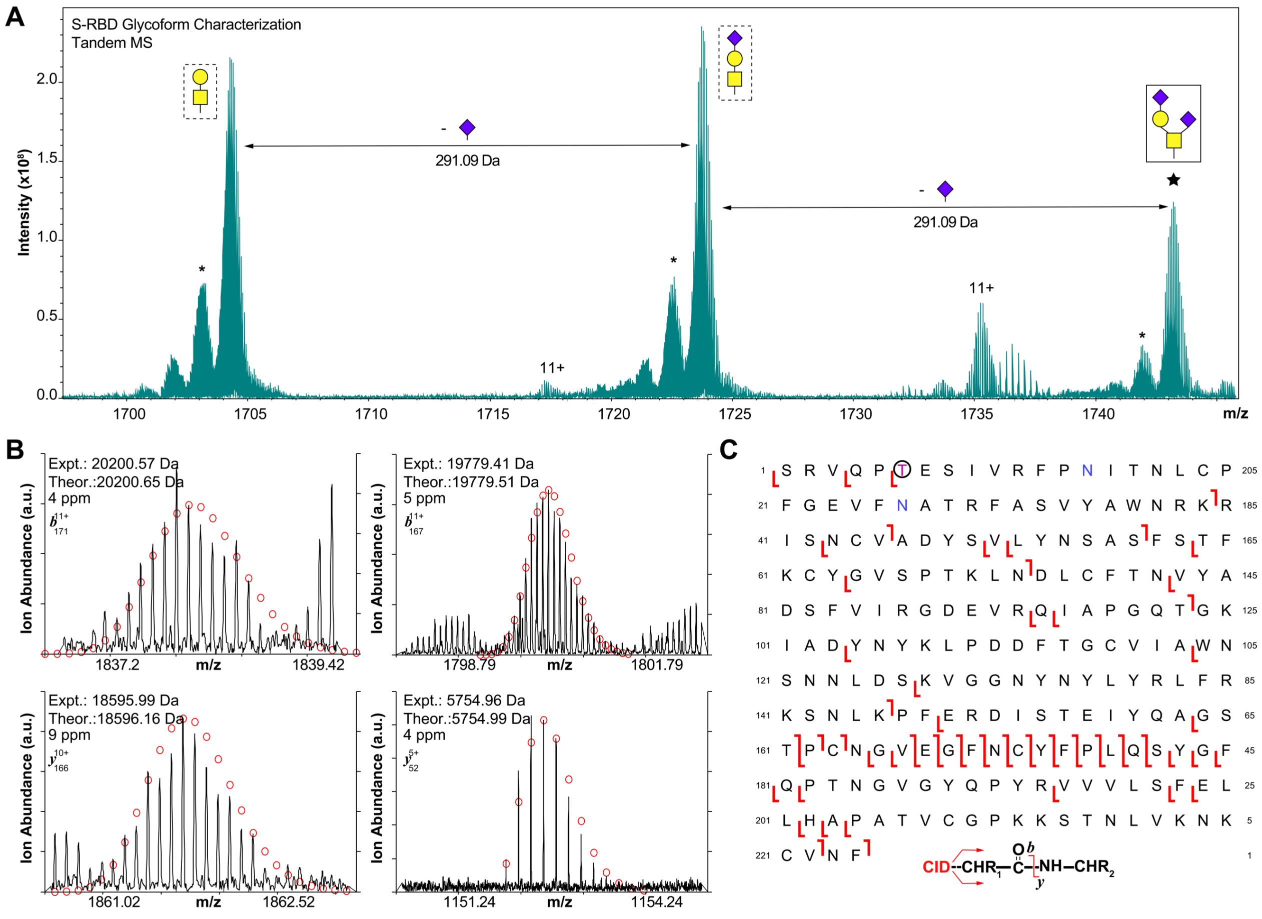


**Figure S8.** (A) MS/MS characterization of S-RBD proteoform isolated from quadrupole window centered at 1743.2 *m/z*, corresponding to the MS[1] shown in Fig. 5. Glycoform characterization reveals the specific S-RBD proteoform to have Core 1 type GalNAcGal(NeuAc)$_2$ glycan. Neutral loss glycan products are labeled. The star represents the 15+ charge state precursor ion and the asterisk "*" denotes an oxonium ion loss. (B) Representative CID fragment ions ($y_{224}^{15+}$, $y_{57}^{4+}$, $b_{166}^{15+}$, and $y_{52}^{4+}$) obtained from S-RBD with glycosite at Thr323. Theoretical ion distributions are indicated by the red dots and mass accuracy errors are listed for each fragment ion. (C) Fragmentation mapping of the specific S-RBD proteoform. Amino acid sequence (Arg319-Phe541) was based on the entry name P0DTC2 (SPIKE_SARS2) obtained from the UniProtKB sequence database. N-terminal Ser is due to removal of signal peptide tPA following cell expression. The blue N (Asn) denotes deamidation following PNGase F treatment.

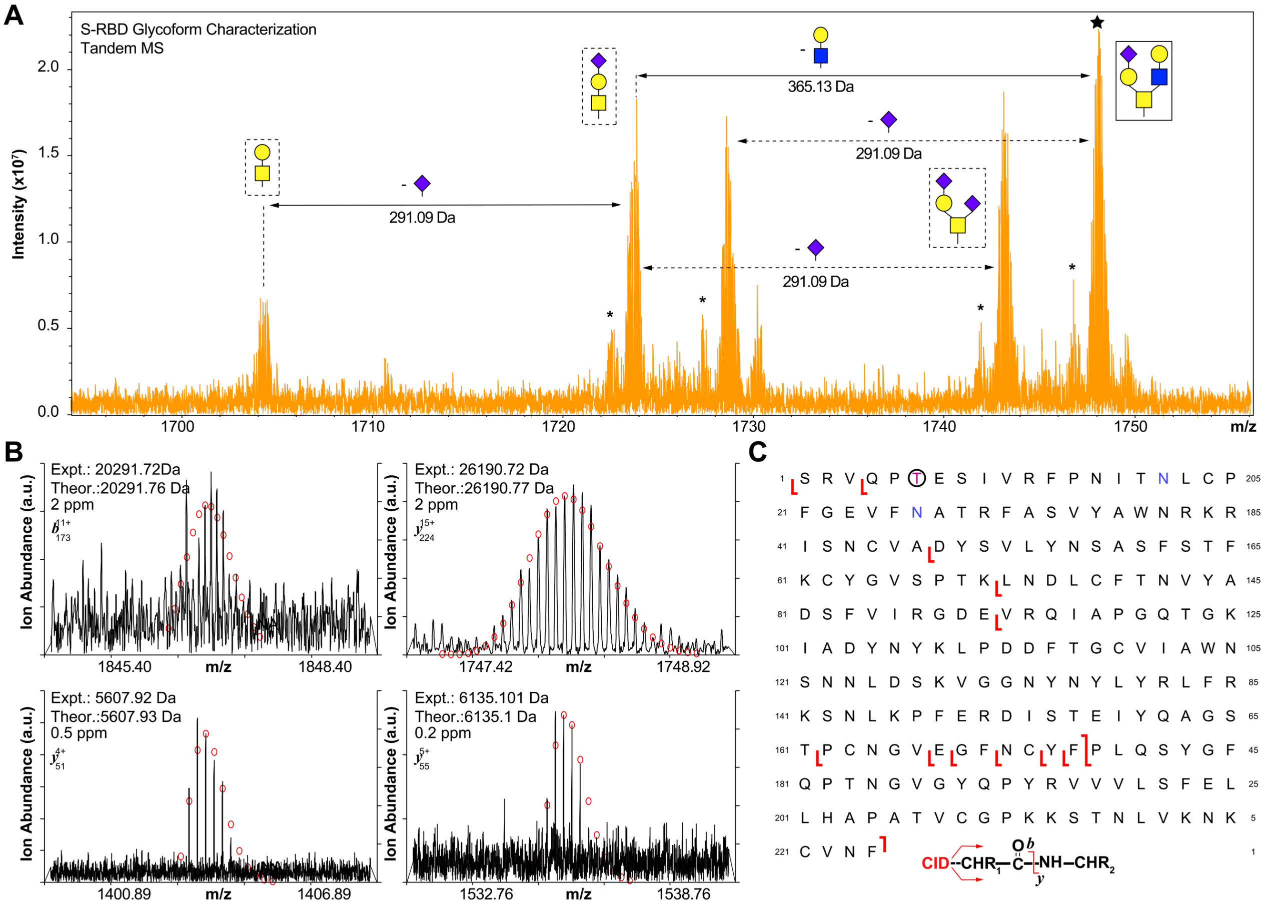


**Figure S9.** (A) MS/MS characterization of S-RBD proteoform isolated from quadrupole window centered at 1748.1 *m/z*, corresponding to the $MS^1$ shown in Fig. 5. Glycoform characterization reveals the specific S-RBD proteoform to have have Core 2 type GalNAcGal(GalNeuAc)(GlcNAcGal) glycan. Neutral loss glycan products are labeled. The star represents the 15+ charge state precursor ion and the asterisk "*" denotes an oxonium ion loss. (B) Representative CID fragment ions ($b_{173}^{11+}$, $y_{224}^{15+}$, $y_{51}^{4+}$, and $y_{55}^{5+}$) obtained from S-RBD with glycosite at Thr323. Theoretical ion distributions are indicated by the red dots and mass accuracy errors are listed for each fragment ion. (C) Fragmentation mapping of the specific S-RBD proteoform. Amino acid sequence (Arg319-Phe541) was based on the entry name P0DTC2 (SPIKE_SARS2) obtained from the UniProtKB sequence database. N-terminal Ser is due to removal of signal peptide tPA following cell expression. The blue N (Asn) denotes deamidation following PNGase F treatment.

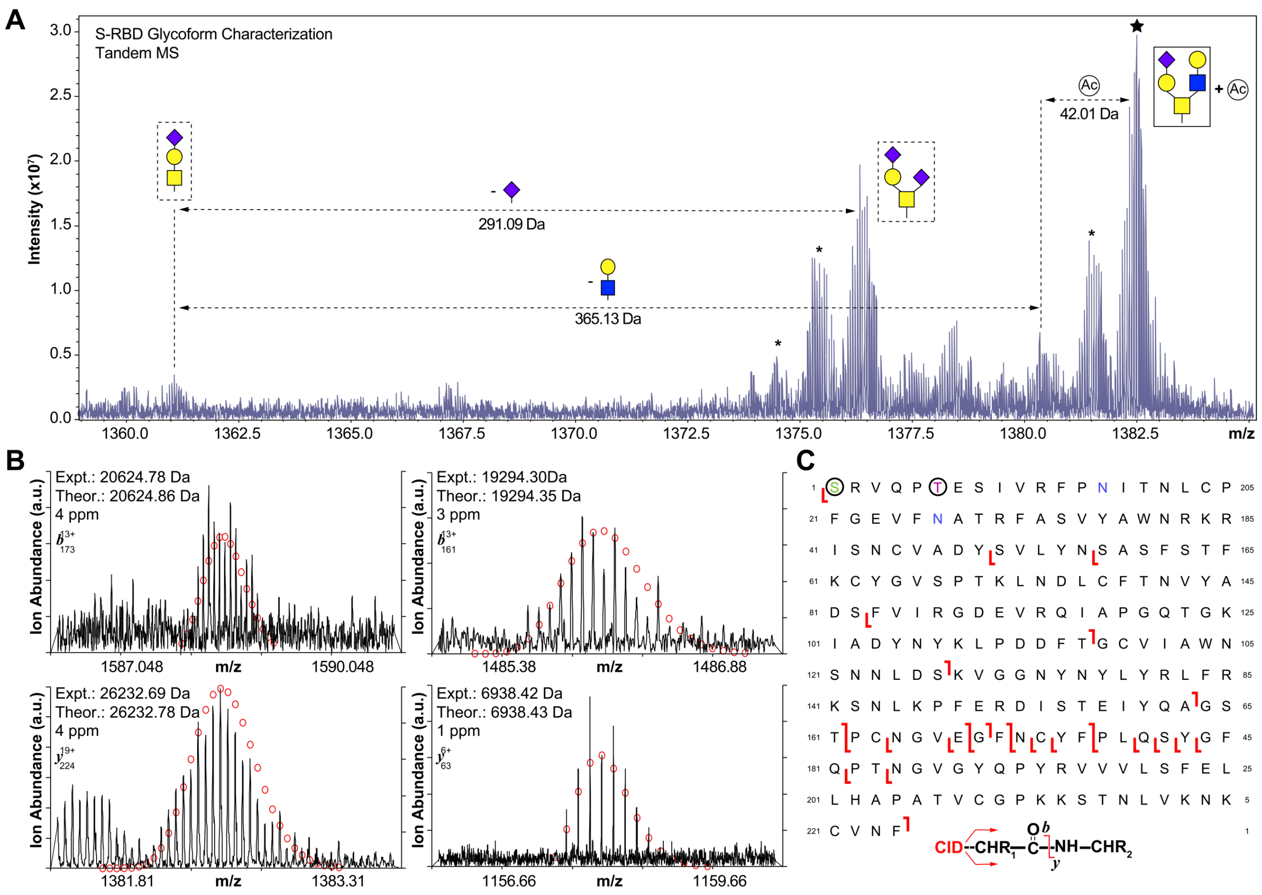


**Figure S10.** (A) MS/MS characterization of S-RBD proteoform isolated from quadrupole window centered at 1382.7. *m/z*, corresponding to the MS[1] shown in Fig. 5. Glycoform characterization reveals the specific S-RBD proteoform to have Core 2 type GalNAcGal(GalNeuAc)(GlcNAcGal) glycan. Neutral loss glycan products are labeled. N-terminal acetylation (Ac) is labeled and corresponds to a +42 Da mass shift. The star represents the 19+ charge state precursor ion and the asterisk "*" denotes an oxonium ion loss. (B) Representative CID fragment ions ($b_{173}{}^{13+}$, $b_{161}{}^{13+}$, $y_{224}{}^{19+}$, and $y_{63}{}^{6+}$) obtained from S-RBD with glycosite at Thr323. Theoretical ion distributions are indicated by the red dots and mass accuracy errors are listed for each fragment ion. (C) Fragmentation mapping of the specific S-RBD proteoform. Amino acid sequence (Arg319-Phe541) was based on the entry name P0DTC2 (SPIKE_SARS2) obtained from the UniProtKB sequence database. N-terminal Ser is due to removal of signal peptide tPA following cell expression. The blue N (Asn) denotes deamidation following PNGase F treatment.

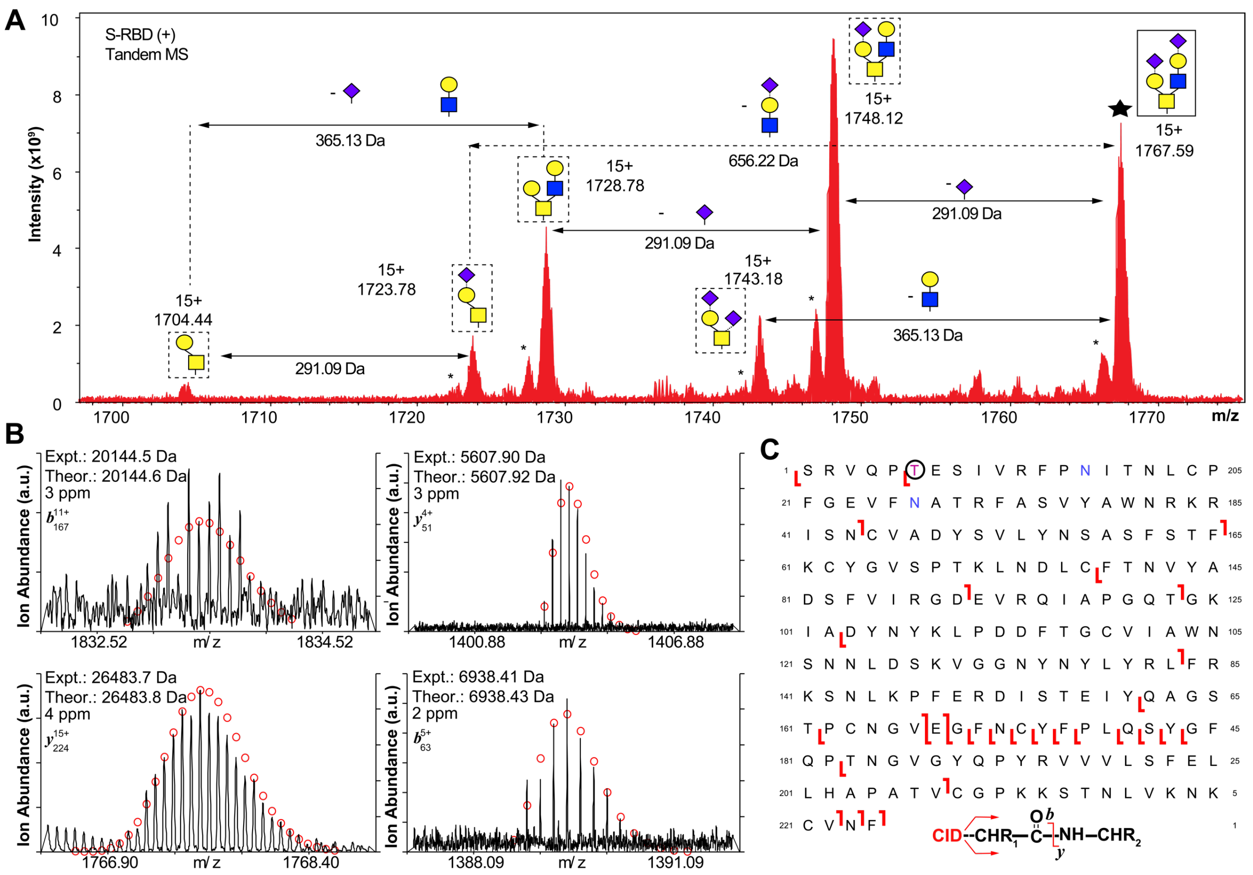


**Figure S11.** (A) MS/MS characterization of S-RBD proteoform isolated from quadrupole window centered at 1767.6 *m/z*, corresponding to the $MS^1$ shown in Fig. 5. Glycoform characterization reveals the specific S-RBD proteoform to have Core 2 type GalNAcGal(GalNeuAc)(GlcNAcGalNeuAc) glycan. Neutral loss glycan products are labeled. The solid star represents the 15+ charge state precursor ion corresponding to 1767.6 *m/z*. The asterisk "*" denotes an oxonium ion loss. (B) Representative CID and ECD fragment ions ($b_{167}^{11+}$, $c_{224}^{15+}$, $y_{225}^{5+}$, and $b_{63}^{5+}$) obtained from S-RBD proteoform isolated from 1767.6 *m/z* precursor with glycosite at Thr323. Theoretical ion distributions are indicated by the red dots and mass accuracy errors are listed for each fragment ion. (C) Fragmentation mapping of the specific S-RBD proteoform corresponding to (B). Amino acid sequence (Arg319-Phe541) was based on the entry name P0DTC2 (SPIKE_SARS2) obtained from the UniProtKB sequence database. N-terminal Ser is due to removal of signal peptide tPA following cell expression. The blue N (Asn) denotes deamidation following PNGase F treatment.
